# Beyond single-trait GxE: higher-order environmental interactions and clonal diversity govern trait relationships in yeast

**DOI:** 10.1101/2025.08.21.671491

**Authors:** Andrea Pozzi, Destdina Korkmaz, Jochen B. W. Wolf

## Abstract

Predicting organismal responses to complex ecological change requires understanding how multiple environmental stressors (E) and genetic background (G) interact on phenotypic variation (P). Here, we investigate how temperature (E_T_) and salinity (E_S_) shape growth (P_G_) and flocculation (P_F_) in multiple strains of the fission yeast, *Schizosaccharomyces pombe*. We find that environmental interactions are critical, with the effect of temperature on flocculation being inverted by changes in salinity (P∼E×E). Multi-stress reaction norms are genotype-dependent (P∼G×E×E), revealing that evolution can adopt diverse strategies to deal with E×E interactions. We further show that the covariation between traits (P×P) is itself a plastic and evolvable feature. The relationship between growth and flocculation changed from negative to positive or completely uncoupled, depending on the specific G×E×E context. This context-dependent covariation suggests that clonal variability and stochastic processes modulate phenotypic outcomes beyond deterministic genotypic effects. Our findings demonstrate that higher-order interactions govern not only individual traits but also their interrelationships, highlighting the necessity of integrating G×E×E effects and phenotypic covariation into models of adaptation.

## Introduction

Organisms inhabit dynamic environments with conditions shifting across spatial and temporal scales. Some of these shifts are gradual, allowing populations to adapt through evolutionary change over generations. Others are abrupt, demanding more rapid phenotypic responses. To cope with such environmental variability, organisms rely on genetic adaptation and phenotypic plasticity (Bradshaw, 1965; Price *et al*., 2003; Bonamour *et al*., 2019), and the interaction between the two: phenotypic plasticity can reduce evolutionary potential (Oostra *et al*., 2018), or conversely, facilitate adaptation to novel stressors (Coates *et al*., 2025). When individuals respond to environmental variation through phenotypic plasticity (P ∼ E), they adjust trait values within the limits of their genetically encoded reaction norms (Fox *et al*., 2019). Long-term evolutionary change requires mutations that alter heritable trait means and variances or introduce novel traits entirely (genotype–phenotype map P ∼ G). Mutations may also introduce variation to reaction norms such that in genetically diverse populations, different genotypes may respond differently to the same environmental stressor (genotype-by-environment interactions P ∼ G×E; (Via & Lande, 1985). These interactions correspond to variation in plasticity, which may itself be heritable and subject to selection (Nussey *et al*., 2005). G×E interactions are widely recognized as important evolutionary drivers, and have been extensively studied, particularly in response to well-characterized environmental factors such as temperature. For instance, research on fruit flies (*Drosophila*) has shown that mitochondrial reaction norms correspond to the thermal conditions of source populations (Camus *et al*., 2017; Lajbner *et al*., 2017). Flies with mitochondria derived from warmer environments recover faster from heat-induced stress, illustrating how genetic variation interacts with environmental pressures to shape physiological responses.

In nature, phenotypic responses to environmental changes are further complicated because they rarely occur in isolation. Instead, organisms are simultaneously exposed to multiple environmental stressors (E×E), such that phenotypic expression is generally shaped by the combined influence of genotype and interacting environmental variables (P ∼ G×E×E; (Via & Lande, 1985; Hietpas *et al*., 2013; Camus *et al*., 2017; Schmidlin *et al*., 2024). These interactions remain relatively understudied, primarily due to the required complexity of experimental design (Schmidlin *et al*., 2024). Yet, it is well conceivable that they are fundamentally shaping fitness landscapes across a large range of organisms and ecosystems. For instance, in marine ecosystems, both temperature and salinity influence species distributions (Hastings *et al*., 2020). A study on the marine copepod *Acartia tonsa*, for instance, revealed that ocean acidification unexpectedly buffered the negative effects of warming on reproductive output, demonstrating that two stressors can have antagonistic effects (Garzke *et al*., 2016; Wang *et al*., 2018). Genotype multi-environment interactions even extend to a medical context, as bacterial resistance to antibiotic stress can be altered by temperature regime (Hietpas *et al*., 2013). To summarize, phenotypic variation is shaped by genotype-environment interactions that can range from simple, direct effects of genotype or environment to complex, multi-way dependencies where the combined effect of multiple stressors is unique to each genotype (**Fig. 1B**).

**Figure 1.**
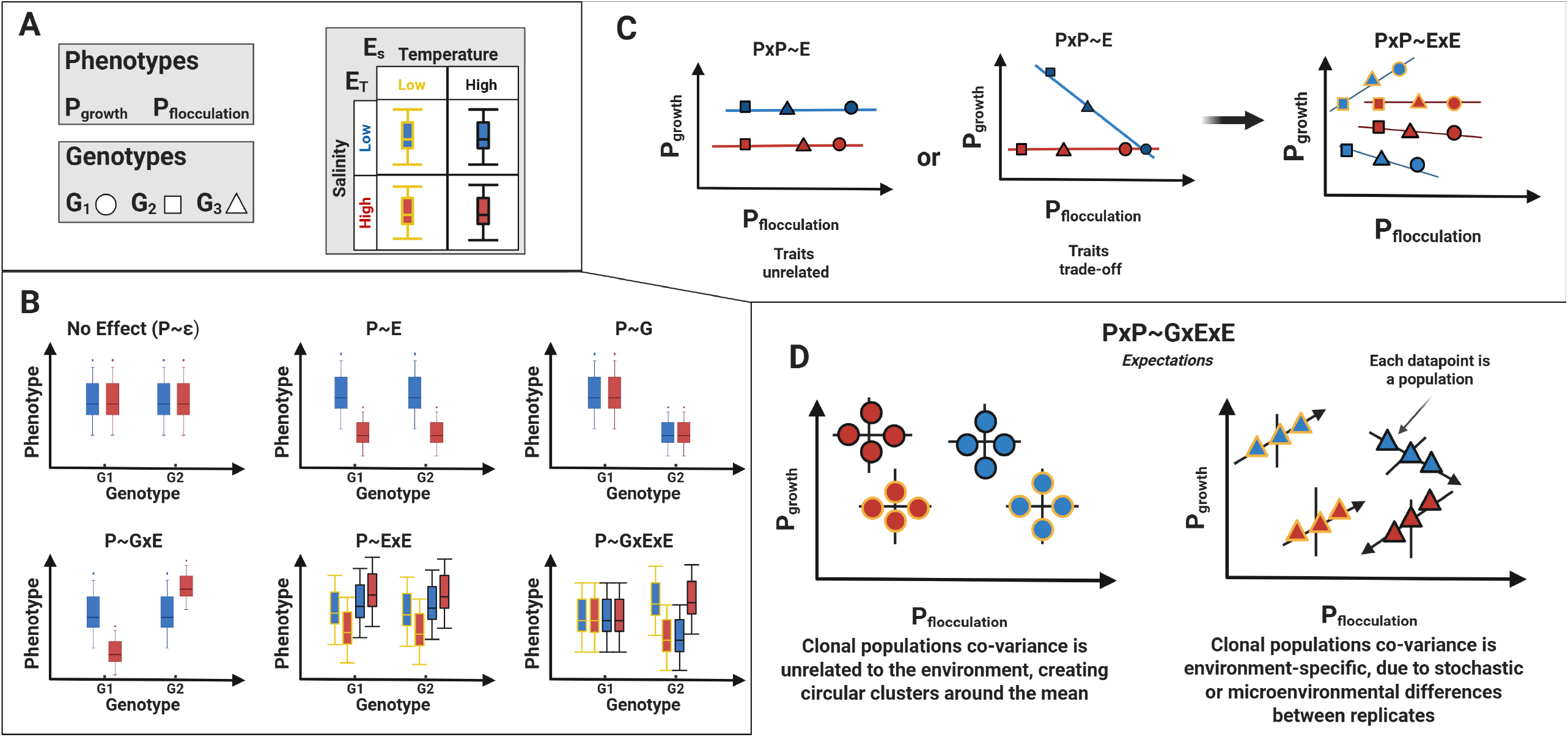
Conceptual framework for G×E interactions and phenotypic covariation. (**A**) *Legend defining the key components*. Three genotypes (G_1_-G_3_) are represented by different shapes. Two phenotypes, growth (P_G_) and flocculation (P_F_), are measured across four environments defined by a factorial combination of salinity (E_T_: Low = blue box, High = red box) and temperature (E_S_: Low = yellow outline, High = black outline). (**B**) *Six models of phenotypic variation illustrated with boxplots*. The models show how phenotypic variation (P) can be attributed to random (e.g. measurement) error (ε), a main effect of Environment (E), a main effect of Genotype (G), to the interaction between Genotype and Environment (G×E), the interaction between two Environments (E×E), or all three (G×E×E). (**C**) *Models of phenotypic covariation (P×P)*. Genotypes may show differences in trait means, but the relationship between phenotypes can be unrelated to genetic variation (left) or exhibit genetic correlations (middle, right). In the example in the middle, genotype 2 has high performance for growth, but low performance for flocculation under low salinity. Genotype 1 shows the opposite pattern, genotype 3 is intermediate resulting in a negative correlation (or trade-off). Under high salinity there is no relationship (P×P∼E). Genetic correlations can further be modulated by environmental interactions (E×E, right). (**D**) *Phenotypic covariation can also exist within clonal populations, but may differ by genotype (P×P∼ G×E×E)*. Left: Shown is the traditional expectation for a single genotype (G1). Growth and flocculation phenotypes differ by environment (P_G_ ∼ E_S;_ P_F_ ∼ E_T_). For repeated measures of clonal populations phenotypes are uncorrelated within each environmental condition (circular clusters). Variation between replicates is due to measurement error or other factors not accounted for. Right: Another genotype (G3) exhibits a different covariation structure within each environment. Phenotypic measurements between replicates show environment-specific correlations between the two traits which may arise by covariation of measurement error or biologically relevant, stochastic physiological or microenvironmental differences between clonal populations.

In addition, it does often not suffice to consider phenotypic traits in isolation. Many traits evolve in concert, constrained by trade-offs or other forms of genetic correlation. The trade-off between growth and reproductive output is a key example: a genotype can either maximize reproduction or growth, but hardly both (Tusso *et al*., 2021). Genetic covariance structure underlying phenotypic correlations thus restricts the degree of freedom for the evolutionary trajectory of a population (Schluter, 1996). Selection will more readily push a population along the ‘path of least resistance’ dictated by genetic correlations, constraining evolutionary change, as because moving outside of these innate genetic correlations is more difficult (Wilke, 2015; Ungar & Hlusko, 2016). The very nature of these trait correlations (P×P) can, in addition, be context-dependent, shifting with environmental conditions (**Fig. 1C**). Importantly, phenotypic correlations need not merely be visible across genetic variation within or between populations. They may also emerge between genetically identical cells, or cell populations, depending on their physiological state (**Fig. 1D**). Such clonal variability, or variation among genetically identical individuals, can be a crucial adaptive strategy. In isogenic populations of the green alga *Chlamydomonas reinhardtii*, for example, individual cells exhibit significant heterogeneity in their growth rates and accumulation of storage compounds (Collins *et al*., 2006; Damodaran *et al*., 2015). This pre-existing variation can act as a bet-hedging mechanism, ensuring that when faced with sudden environmental stress like nutrient starvation, a subpopulation of cells is already primed to survive, thereby enhancing population resilience without genetic change.

Despite growing efforts towards more complex experimental designs addressing genotype-environment interactions, most studies still tend to focus on single traits or environmental variables (Hoffmann & Sgrò, 2011; Lajbner *et al*., 2018; Tusso *et al*., 2021), overlooking the complex, often counter-intuitive nature of these high-order interactions. Microorganisms are particularly well-suited for experimentally dissecting these complex interactions, from G×E×E dependencies to the dynamics of clonal variability (Merritt, 1966; Kaviriri *et al*., 2020). Their short generation times and ease of replication allow for the direct observation of evolutionary processes across many generations, which is impractical in long-lived organisms. Furthermore, their predominantly clonal reproduction provides a powerful framework for studying phenotypic plasticity in traits and trait co-variance with full control over genetic variation. Following this rationale, we used the unicellular fungus *Schizosaccharomyces pombe* as a model system to investigate how simultaneous changes in temperature and salinity (E_T_, E_S_) influence two key fitness-related traits, growth and flocculation (P_G_, P_F_). We employ a full factorial design with nine haploid strains of varying genetic distance (G), from closely-related sibling strains to divergent natural accessions (Westermann *et al*., 2022). Measuring two traits across 2944 population allows us to characterize not only how different genotypes express traits under interacting stressors, but also how this G × E × E context shapes the phenotypic covariation (P × P)among clonal cell populations.

## Results

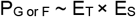

We first explored phenotypic plasticity for growth (P_G_) and flocculation (P_F_) separately across both environments using the full factorial design P_G or F_ ∼ E_T_ × E_S_. To minimize genotypic variation, we initially used six near-clonal genotypes derived from a population of strain JB900 (differing by <500 SNPs, **Fig. S1**) for experimental validation (**Fig. 2A, B**). For growth, temperature reaction norms showed only a modest interaction with salinity which was significant for all of the six strains (F_3,312_ > 6.44, p < 0.0003). Some strains like JB900_6 showed slightly better growth at high temperature under a low than under high salinity regime. Flocculation showed a strong and pervasive interaction across all six genotypes (F_3,312_ > 31.93, p < 2 × 10^-16^): flocculation under low salinity increased with temperature, peaking at 38 °C. This pattern was reversed under high salinity, where flocculation was highest at 29°C and decreased as temperature rose. These findings indicate different dual-stress reaction norms for the two phenotypes, with response patterns being consistent across the near-clonal genotypes.

**Figure 2.**
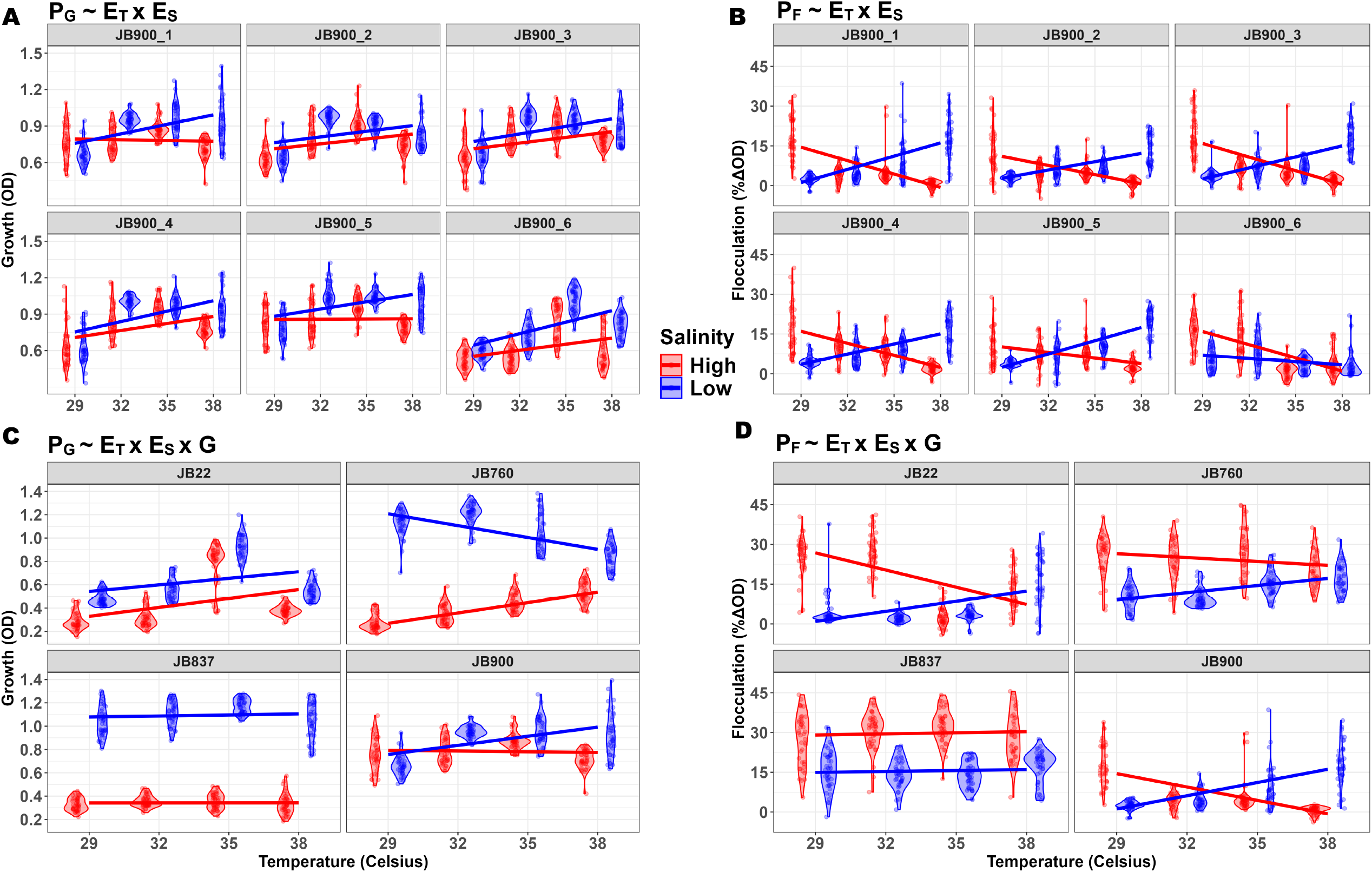
G×E×E interactions for growth and flocculation in yeast (P_G_ or P_F_). Phenotypic responses of different yeast strains to a factorial combination of temperature (x-axis) and salinity (Low = blue, High = red). Each plot displays raw data as jittered points representing a minimum of 40 replicates, the data distribution as violin plots, and the overall trend as a linear regression line fitted separately for each strain and salinity regime. **(A, B)** Growth (OD) and flocculation (%ΔOD) responses for six low-diversity yeast strains modeled as a function of the interaction between temperature (E_T_) and salinity (E_S_). **(C, D)** The same interactions are shown in four high-diversity strains (JB22, JB768, JB837, JB900). These panels illustrate how phenotypic plasticity is affected by the interaction between environmental variables and the specific genotype.

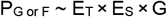

We then extended the analysis to include genotypic effects (G) (**Fig. 2C - D**), adding three genetically divergent strains. For both growth and flocculation, the interaction of salinity and growth reaction norms depended on the genetic background of the strain (P_G_: F_3,1312_ = 67.63 and P_F_: F_3,1312_ = 51.06, p < 2 × 10^-16^ for both, see **Tab. S1** for all measurements and **Tab.S2** for statistical result of the full three-way interaction and strain-specific two-way analyses). **Figure 2C** displays the change in growth performance which was governed mainly by salinity, and strain-specific interaction with temperature: JB22 and JB900 showed a temperature-dependent growth response without evidence of an interaction with salinity. By contrast, JB837 showed a strong, yet temperature-independent reduction in growth at high salinity indicating pervasive low salinity tolerance. JB760 also grew substantially worse at high salinity, but sensitivity to salinity was temperature-dependent (E_T_ × E_S_): growth performance decreased at low salinity but increased at high salinity. For flocculation dual-stressor reaction norms were also strain specific (**Fig. 2D**): in JB22, JB760 and JB900 flocculation was favored by high temperatures under low salinity, under high salinity the pattern was reversed to different degrees. The E_T_ × E_S_ interaction was marked in all three strains, and most pronounced for JB900 (cf. **Fig. 2B**). In contrast, JB837, showed the weakest E_T×_E_S_ interaction, with high salinity consistently elevating flocculation across temperatures. These differences highlight the complexity of E×E interactions and demonstrate that genetic background plays a crucial role in shaping plastic responses of yeast strains to multiple environmental factors acting at the same time.

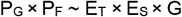

We next examined the effect of genotype and environment on the covariation between growth and flocculation (P_G_ × P_F_ ∼ E_T_ × E_S_ × G), which is conceptually similar to a multivariate reaction norm (**Fig. 3**). Note, however, that in this experiment phenotypic covariation (P_G_ × P_F_) between replicate populations within each strain cannot be due to genetic variation, since we expect far less mutations within a clonal population than between JB900 variants showing consistent phenotypes (see **Fig. 2A, B**). Within each strain, phenotypic covariation among clonal populations can thus only arise from correlated measurement error, intrinsic dependencies of the metrics, or biologically meaningful, stochastic physiological or microenvironmental differences. While the former sources would produce covariation that is consistent in direction and strength, biologically induced covariation may vary depending on the environmental and genotypic context (see **Fig. 1C, D**).

**Figure 3.**
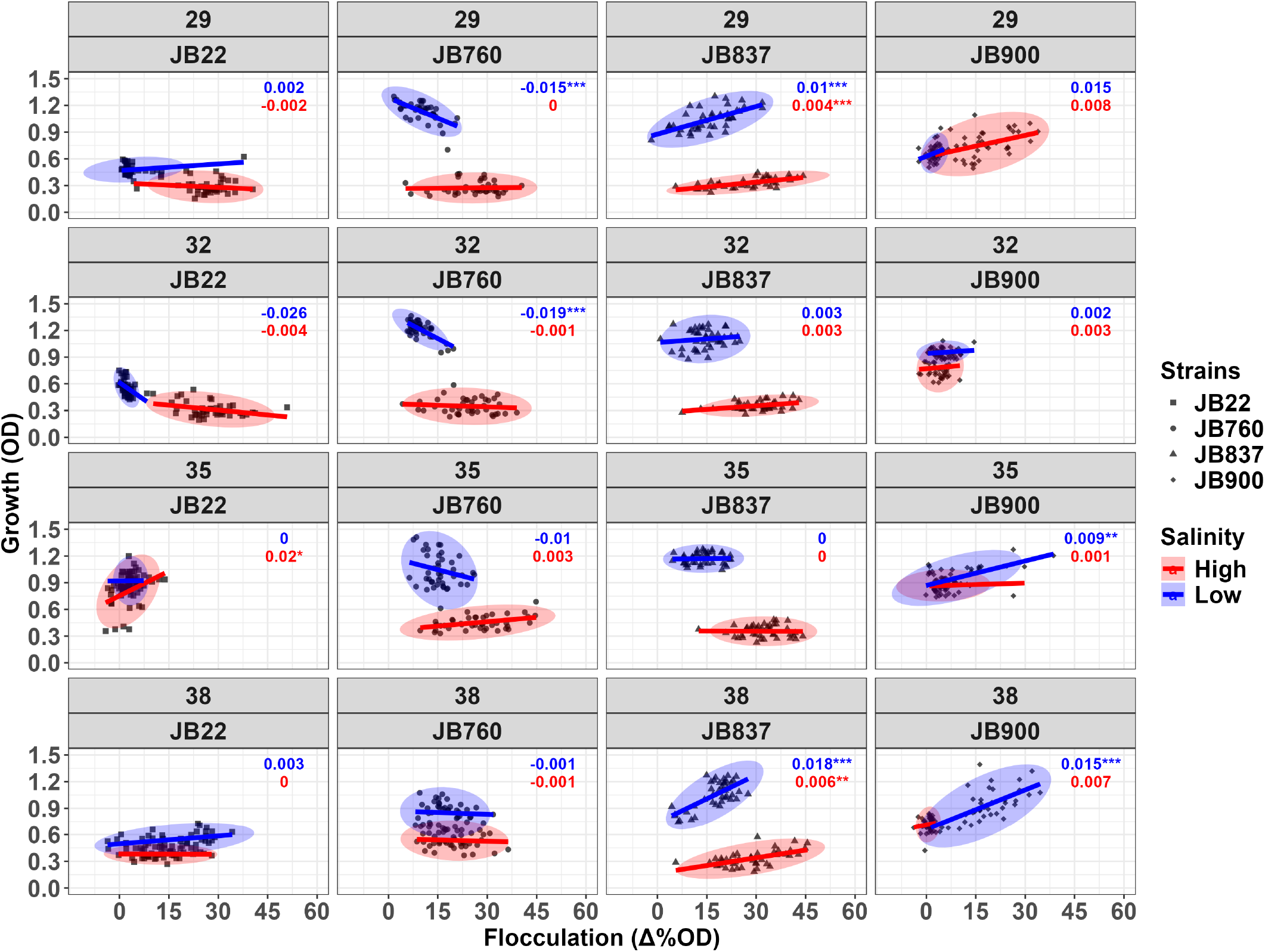
G×E×E interactions on phenotypic covariation (P_G ×_ P_F_). Each panel shows the relationship between growth (y-axis) and flocculation (x-axis) for a unique combination of yeast strain (columns) and temperature (rows). Within each panel, data from high salinity conditions are shown in red and low salinity in blue. Linear regression lines and ellipses illustrate the correlation between the two traits, with inset numbers indicating Pearson correlations for each set of ≥ 40 clonal population replicates; asterisks denote significance (*p* < 0.05, \**p* < 0.01, \*\**p* < 0.001) for Bonferroni-corrected p-values. The figure demonstrates that the relationship between growth and flocculation is highly plastic, shifting from negative correlation (e.g., in strain JB760) to a positive correlation (e.g., in strains JB837 and JB900) depending on the specific genotype and environmental context.

Covariation between phenotypic traits was influenced both by environmental context and strain. For instance, for JB22 growth and flocculation were largely independent across almost all conditions. In contrast, strain JB760 exhibited a conditional trade-off: at 29°C and 32°C flocculation on growth showed a negative correlation (r = −0.015, adj-p < 0.005) which disappeared at higher temperatures. Other strains displayed positive relationships. Strain JB837 had a general positive relationship under low salt conditions, which was most visible under cold and heat stress at 29°C and 38°C, respectively (r = 0.01, 0.018, adj-p < 0.005). Strain JB900 presented an even stronger, positive relationship between flocculation and growth under low salt intensifying with heat stress (r = 0.015, adj-p < 0.005 at 38°C). This context-dependent covariation is unlikely to be a mere artifact. Multivariate Analysis of Variance with growth and flocculation as combined phenotypic response variables, revealed statistical significance for the critical three-way interaction between salinity, temperature, and strain (F_24,3248_ = 29.4, p < 2.2×10^−16^, see **Tab. S2**). We therefore interpret this context-dependent covariation as a biological signal rather than a technical artifact.

## Discussion

This study suggests that predicting phenotypic outcomes in multi-stressor environments requires moving beyond simple additive models of environmental and genetic effects. Our first key finding reveals that plasticity in trait expression is not adequately captured by single-factor reactions norms. Instead, it critically depends on the specific interaction between environmental factors (P_G or F_ ∼ E_T_ × E_S_). Moreover, the importance of E × E interaction varied by trait, with a pervasive interaction effect on flocculation, in contrast to a more modest interaction observed for growth. The consistency of the dual-environment reaction norms across near-clonal JB900 replicates validates the experimental control and establishes these complex interactions as a genuine biological phenomenon.

The introduction of genetic diversity in the form of divergent strains revealed that these dual-environment reaction norms are also dependent on the genotype (P_G or F_ ∼ E_T_ × E_S_ × G). For example, in JB837, both phenotypes were strongly affected by salinity, but were indifferent to temperature. JB22 and JB900 were minimally affected by any environmental factor for growth, but showed a marked crossover E_T_ × E_S_ interaction for flocculation. G × E × E interactions provide the raw material for the evolution of multi-environment plasticity, but it also fundamentally limits our ability to predict evolutionary trajectories. For example, global warming causes multiple environmental variables to change simultaneously (e.g. ocean temperature and salinity). The resulting “lag” between a population’s traits and the new environmental optimum might, to some degree, be buffered by the ‘correct’ plastic response to both stressors. Our results show that the reaction norms in such a situation depend on the underlying genotype. Provided enough genetic variation, we therefore expect selection on genotypes carrying the right combination of plastic responses to help populations adapt to multiple, simultaneous stressors. While our results suggests that there is scope for evolution to shape adaptive plasticity, the underlying complexity makes it exceedingly difficult to identify future evolutionary “winners and losers” under climate change (Hoffmann & Sgrò, 2011). Yet, maintenance of genetic variation appears to be key to maintain a reservoir of adaptive, multi-environmental responses allowing populations to persist.

Perhaps our most puzzling finding is that the covariation between traits was itself environmentally plastic and genotype-dependent (P × P ∼ E × E × G). Clonal populations showed an environment- and genotype-dependent relationship between growth and flocculation: a temperature-sensitive negative relationship in JB760, a stress-activated positive relationship in JB837 and JB900, and lack of correlation in JB22. This context-dependent covariation, suggests that we observe a biologically meaningful signal rather than a technical artifact. Trait covariation between clonal cells, or cell populations, has been rarely reported (Levy *et al*., 2012), but points to the importance of ‘clonal variability’ (Spudich & Koshland, 1976; Lidstrom & Konopka, 2010; Sherry & Rego, 2024), where stochastic physiological or microenvironmental differences among genetically identical populations lead to divergent phenotypic outcomes. This phenomenon challenges traditional adaptation models by showing that even with genotype and macro-environment held constant, phenotypic solutions can vary. This implies that evolutionary trajectories might not only be affected by genetic correlations, but additionally by gene-environment interactions modulating the mode and strength of plastic responses within clonal cell populations.

In summary, by demonstrating that higher-order interactions govern not only individual traits but also the relationships between them, our findings underscore the need to integrate G×E×E effects and phenotypic covariation into evolutionary models. For microbial populations, where rapid reproduction can amplify the effects of stochasticity and microenvironmental heterogeneity, understanding these complex, multi-layered interactions is essential for understanding adaptation in a changing world. The results may similarly be of relevance to medical application, for example in semi-clonal cancer cell populations exposed to multi-medication environments.

## Materials and Methods

### Yeast strains

The six low-diversity strains of fission yeast (*S. pombe*) used in this study were generated by crossing two parental strains, themselves derived from a single yeast population (JB900), which had fewer than 1000 unique SNPs. The resulting offspring differed by less than 500 unique SNPs (**Fig. S1**) corresponding to an average genetic distance (*d*) of 0.008 (**Tab. S3**), calculated using TASSEL (Bradbury *et al*., 2007). The four divergent strains used were JB22, JB760, JB837, and JB900_1. Strains JB22 and JB760 are genetically similar (*d* = 0.03) and share a similar ancestry but are distinct from the other two strains (0.3 < *d* < 0.4). JB837 and JB900 are also genetically distinct from each other (*d* = 0.2). The mitochondrial DNA (mtDNA) of all strains also showed high diversity (0.4 < *d* < 0.6) but matching the pattern seen in the nucleus. This suggests is unlikely that the effects E × E × G are driven by the mitochondrial mutations, as seen in other studies on animals (Lajbner *et al*., 2018; Pozzi & Dowling, 2021). Strains were not selected for any specific phenotype and preserved at 4°C. Before each experiment, strains were grown at 32°C in liquid media to saturation, and we took only 200 µL to be tested at specific temperatures and salinities. All experiments were conducted using yeast extract supplemented (YES) medium with 2% glucose. Salinity was adjusted by adding 10 g/L NaCl. Although pilot experiments were performed with up to 20 g/L NaCl, most strains could not tolerate this higher concentration. Therefore, we used a concentration of 10 g/L NaCl for all subsequent experiments.

### Growth Measurement

Yeast growth can be measured in either solid or liquid media using different techniques. Quantitative measurement on solid media typically involves inoculating an agar plate with standardized amounts of yeast at specific positions, often using automated systems for precision. Colony diameter is then measured via imaging techniques (Jeffares *et al*., 2015). This method offers the advantages of high throughput and standardized results, particularly for well-characterized species in appropriately equipped laboratories. In contrast, using liquid media (this study) is more cost-effective and flexible, as it requires less specialized equipment and is applicable to a broader range of species (Stevenson *et al*., 2016). Growth measurement in liquid media is based on absorbance. Cells are irradiated with light at a specific wavelength (typically 600 nm), and the resulting optical density (OD) is measured. At low cell densities, the OD is proportional to the cell concentration according to the Beer-Lambert law (Myers *et al*., 2013). Here, *A* represents absorbance, *ϵ* is the molar absorptivity, *l* is the optical path length, and c is the concentration (approximating the number of cells).

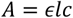

The relationship between OD and absolute fitness is not direct, as the accuracy of OD600 measurements decreases at higher cell densities (Stevenson *et al*., 2016). Consequently, we treated growth as a distinct trait rather than a direct proxy for fitness, since our measurement encompasses both cell proliferation and survival. All OD600 measurements were performed using a Biotek Powerwave XS plate reader (Agilent Technologies), which measures the absorbance in 96-well plates (8×12) by passing light through the bottom of the plate. We opted not to generate growth curves because significant liquid evaporation occurred from temperatures starting at 32°C. Using a plate cover failed to prevent this issue and introduced additional noise to the measurements by limiting oxygen. Therefore, yeast cultures were grown in deep-well plates (1 mL capacity) for 24 hours and then transferred to shallower plates (300 µL capacity) for measurement. This procedure allowed us to take a single endpoint OD600 measurement at 24 hours, which we used as a proxy for total growth. To mitigate the effects of cell sedimentation, cultures were mixed first by pipetting and then by a one-second, low-intensity shake in the plate reader. The growth data are based solely on the OD measurement taken after this initial shake; subsequent shakes and measurements were used to estimate flocculation.

### Flocculation measurement

We adapted the growth measurement protocol to quantify flocculation. The rationale was to measure changes in turbidity via OD as an alternative to using a specialized instrument to determine a flocculation index (Vidgren & Londesborough, 2011; Stewart, 2018). The plate reader illuminates each well of a 96-well, optically clear, flat-bottom plate from below. We observed that while letting a sample stand slightly increased its absorbance, shaking the sample produced a much larger increase. This observation suggests that a higher absorbance reading is obtained when yeast flocs are dispersed. Therefore, a sample whose absorbance does not increase after shaking is considered non-flocculated. Conversely, an increase in absorbance with each shake indicates the presence of flocs that are being progressively broken apart. To quantify flocculation, we measured the OD600 three consecutive times, with a one-second, low-intensity shake preceding each measurement. We then calculated the change in OD (ΔOD) by subtracting the initial reading from the final reading.

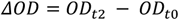

We limited the procedure to three measurements because the OD value typically stabilized by the third reading, suggesting that three shakes were sufficient to fully resuspend the yeast. Because OD600 can be influenced by factors such as cell orientation, the measurement is not perfectly consistent; we observed a few instances of negative ΔOD, likely attributable to such artifacts. A key advantage of this method is its direct comparability to the growth data, as both are derived from OD measurements.

All measures for growth and flocculation were taken for a minimum of 40 replicates (range 40-42) for each of the 8 environmental combinations (2 salinity regimes x 4 temperatures) and in each strain. This resulted in a combined total of 5,888 measurements across nine strains including both growth and flocculation which are provided in **Table S1**.

### Statistical analysis

We performed all statistical analyses using R, primarily using native functions of the base package (“R Core Team, 2021)or from the *ggplot2* package (Wickham, 2016). To test for an environmental interaction, we chose a full factorial design to test for all main effects and interactions.

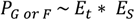

Because the data did not meet the assumptions for standard ANOVA, we used the Aligned Rank Transform (ART) ANOVA, a non-parametric method that allows for robust testing of interactions effects on ranked data (Leys & Schumann, 2010; Wobbrock *et al*., 2011). Two-way Aligned Rank Transform (ART) Analysis of Variance (ANOVA) was performed with the *art()* and *anova()* functions from the *ARTtool* package (Wobbrock *et al*., 2011). Full data about the measurements can be found in **Tab. S1**.

Similarly, to assess the effects of genotypes and environment on single yeast phenotypes, we used a three-way Analysis of Variance (ANOVA) performed with the *aov()* function. For this model as well, we chose a full factorial fixed-effects model to test all main effects and interactions.

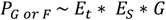

While a mixed-effects model with strain as a random effect could be considered for this experimental design, we opted for a fixed-effects approach. Given the limited number of strains (n = 4), estimating a reliable variance component for a random effect is statistically challenging. Therefore, we treated strain as a fixed factor to directly compare the specific responses among these four genotypes rather than attempting to generalize our findings to a distribution of strains. Following the ANOVA, significant interaction effects were dissected using a Tukey’s HSD post-hoc test using the *TukeyHSD()* function to identify which specific experimental groups differed significantly from one another. The pairwise comparison generated a total of 496 comparisons, and we included them in **Fig. S3-6**).

To understand which factors may influence covariation between phenotypes, we used Multivariate analysis of Variance (MANOVA; Krzanowski, 2000) with growth and flocculation as response, and both environments as explanatory variables.

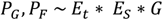

Instead of looking at growth and flocculation in isolation, this analysis examines them as a single, combined phenotype (Yang *et al*., 2010). This analysis treats the phenotype not as two separate numbers, but as a single point on a map where the axes are ‘growth’ and ‘flocculation’. MANOVA then tests if the location of this point systematically shifts when the environment or strain changes (Mérot *et al*., 2016). To explore the significance of results from the MANOVA, we performed a Canonical Variate Analysis (CVA). This analysis identifies the linear combination of the response variables (P_G_ and P_F_) by creating a weighted linear combination of the explanatory variables (G, E_T_ and E_S_) that maximally separates the experimental groups. This is achieved by maximizing the ratio of the variance between the groups to the variance within the groups, a logic identical to the F-statistics in an ANOVA. The resulting canonical variates represent new axes in multivariate space onto which the data can be projected to optimally visualize the significant effects revealed by the MANOVA. Lastly, to analyze the phenotypic covariation between growth and flocculation under each combination of environmental condition (corresponding to **Fig. 3**), we performed a Pearson correlation test using the *cor.test()* function. To account for multiple comparisons across all conditions, the resulting p-values were adjusted using the Bonferroni correction.

## Supporting information

All Supplemental Information

## Acknowledgment

We acknowledge Bart Nieuwenhuis for sharing his expertise on working with *S. pombe* in the laboratory, Dirk Metzler and members of the Division of Evolutionary Biology for fruitful discussion.

## Author contributions

AP conceptualized the work, performed all computational analyses, and wrote the initial draft. DK performed all the measurements and experiments. JW conceptualized the work and wrote the initial draft. All the authors have read and revised the manuscript.

## Supplementary Material Legend

**Figure S1**. Venn diagrams of unique and shared Single Nucleotide Polymorphisms (SNPs) in low- and high-diversity yeast strains. The top panel shows the genetic overlap among the six low-diversity strains (JB900_1 through JB900_6). Numbers in the non-overlapping regions indicate SNPs unique to a single strain (e.g., 188 SNPs are unique to JB900_1), while numbers in overlapping regions represent SNPs shared by two or more strains. All the strains share ∼54k SNPs that differ from the reference strain.

**Figure S2. Diagnostic plots for the normality of model residuals for growth and flocculation**. Top Panel: A histogram of the residuals from the three-way ANOVA for growth is shown with an overlaid kernel density estimate (shaded orange area). The solid blue line represents a theoretical normal curve with the same mean and standard deviation as the growth model residuals. The close alignment between the distribution of residuals and the theoretical curve suggests that the model assumption of residual is normality met for the growth model. Bottom Panel: A histogram of the residuals from the three-way ANOVA for flocculation is shown with an overlaid kernel density estimate (shaded orange area). The solid blue line represents a theoretical normal curve with the same mean and standard deviation as the flocculation model residuals. The distribution exhibits some deviation from the theoretical normal curve, with a slightly more peaked center and potentially heavier tails. However, given the known robustness of ANOVA to moderate deviations from normality, the assumption is likely sufficiently met for the flocculation model as well.

**Figure S3. Pairwise comparisons of environmental effects on growth**. Results of the Tukey’s HSD post-hoc test for the growth phenotype. Each pairwise comparison is a data point, colored simply to better distinguish each comparison, not according to any specific values. The plots show the 95% confidence intervals for the difference in means (Δ mean) for pairwise comparisons. If a confidence interval does not cross the vertical zero line, the difference is statistically significant. Left panel: Comparisons for the main environmental effects (P∼E). Right panel: Comparisons for the two-way environment-by-environment interactions (P∼E×E).

**Figure S4. Pairwise comparisons of environmental effects on flocculation**. Results of the Tukey’s HSD post-hoc test for the flocculation phenotype. Each pairwise comparison is a data point, colored simply to better distinguish each comparison, not according to any specific values. The plots show the 95% confidence intervals for the difference in means (Δ mean) for pairwise comparisons. If a confidence interval does not cross the vertical zero line, the difference is statistically significant. Left panel: Comparisons for the main environmental effects (P∼E). Right panel: Comparisons for the two-way environment-by-environment interactions (P∼E×E).

**Figure S5. Pairwise comparisons of higher-order interaction effects on growth**. Results of the Tukey’s HSD post-hoc test for the growth phenotype, dissecting the Genotype-by-Environment (G×E) and Genotype-by-Environment-by-Environment (G×E×E) interaction terms from the three-way ANOVA. Each pairwise comparison is a data point, colored simply to better distinguish each comparison, not according to any specific values. The plots show the 95% confidence intervals for the difference in means (Δ mean) for specific pairwise comparisons between experimental groups. A confidence interval that does not cross the zero line indicates a statistically significant difference.

**Figure S6. Pairwise comparisons of higher-order interaction effects on flocculation**. Results of the Tukey’s HSD post-hoc test for the flocculation phenotype, dissecting the Genotype-by-Environment (G×E) and Genotype-by-Environment-by-Environment (G×E×E) interaction terms from the three-way ANOVA. Each pairwise comparison is a data point, colored simply to better distinguish each comparison, not according to any specific values. The plots show the 95% confidence intervals for the difference in means (Δ mean) for specific pairwise comparisons between experimental groups. A confidence interval that does not cross the zero line indicates a statistically significant difference.

**Table S1. Final Measurement Data**. The table includes all the final measurements across the Low Diversity (6) and High Diversity strains (3), 2944 measurements for growth and 2944 measurements for flocculation. This does not include repeated measurements to calculate the flocculation.

**Table S2. Statistical Analyses**. The table includes the details for all the statistical interactions described in the manuscript. This includes ANOVA for P∼E×E×G and MANOVA for P×P∼E×E×G interactions using pooled strains. Also to test strain-specific P∼E×E interactions the ART ANOVA was performed for both Low Diversity and High Diversity strains.

**Table S3. Pairwise Genetic Distance**. The table shows the genetic distance from pairwise comparisons of each strain, calculated using TASSEL-I. The three tables show the genetic distance for the whole nuclear genome, for the Chromosome I (the largest across the three chromosomes), and for the mitochondrial DNA.

## Notes

### Competing Interest Statement

The authors have declared no competing interest.

### Summary of Updates

I included the raw data in Table S1. all in the zip file of SI

