## Supplementary figures and images for "Beyond single-trait GxE: higher-order environmental interactions and clonal diversity govern trait relationships in yeast"

### FigS1.png

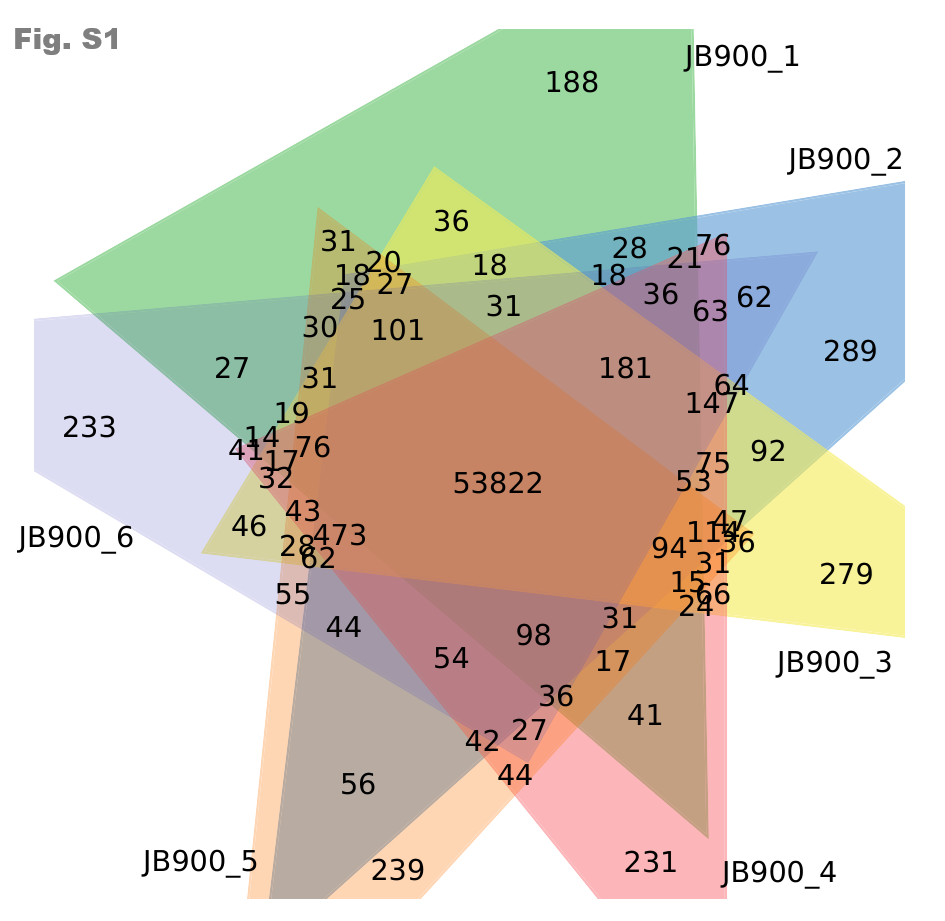

### FigS2.png

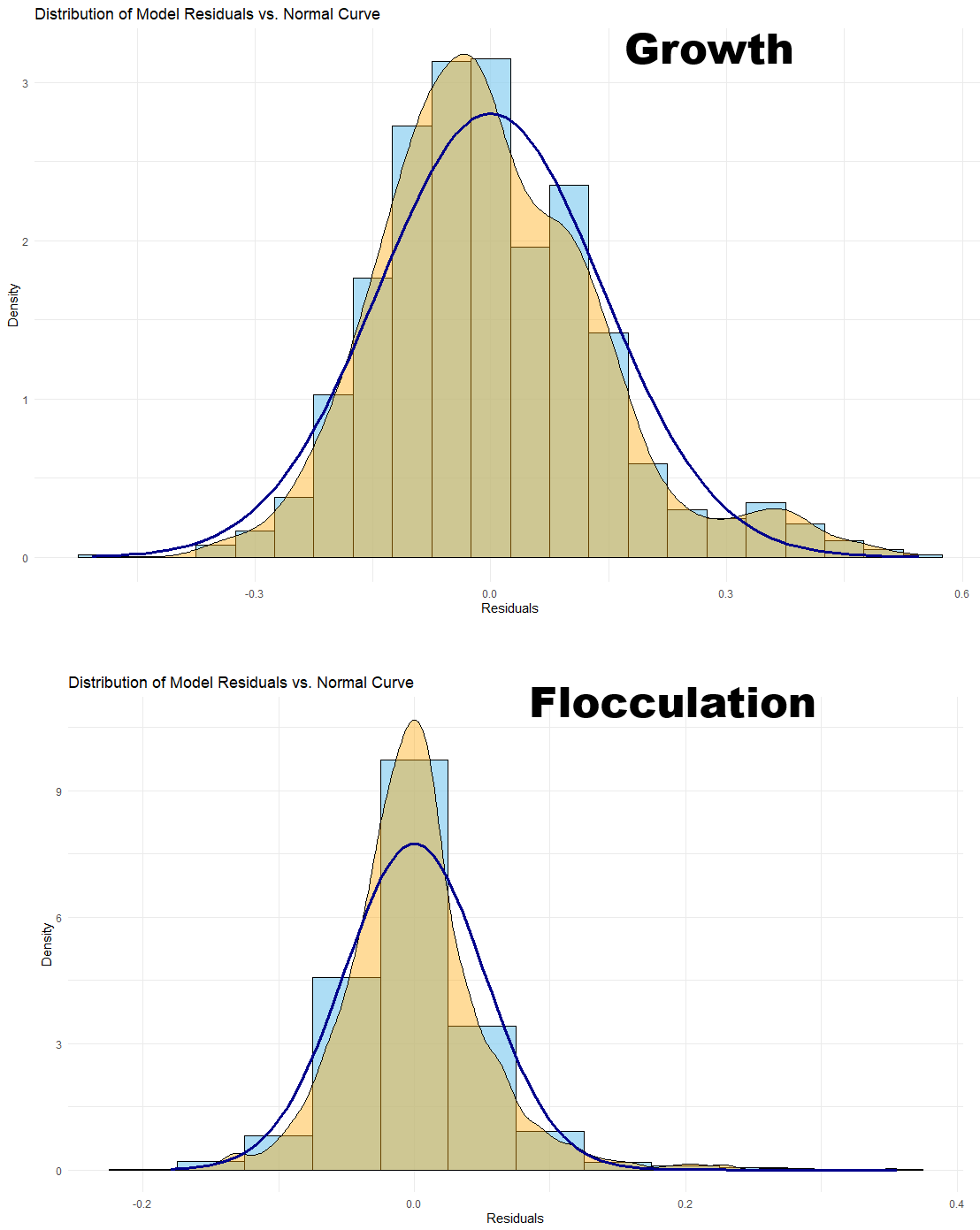

### FigS3.png

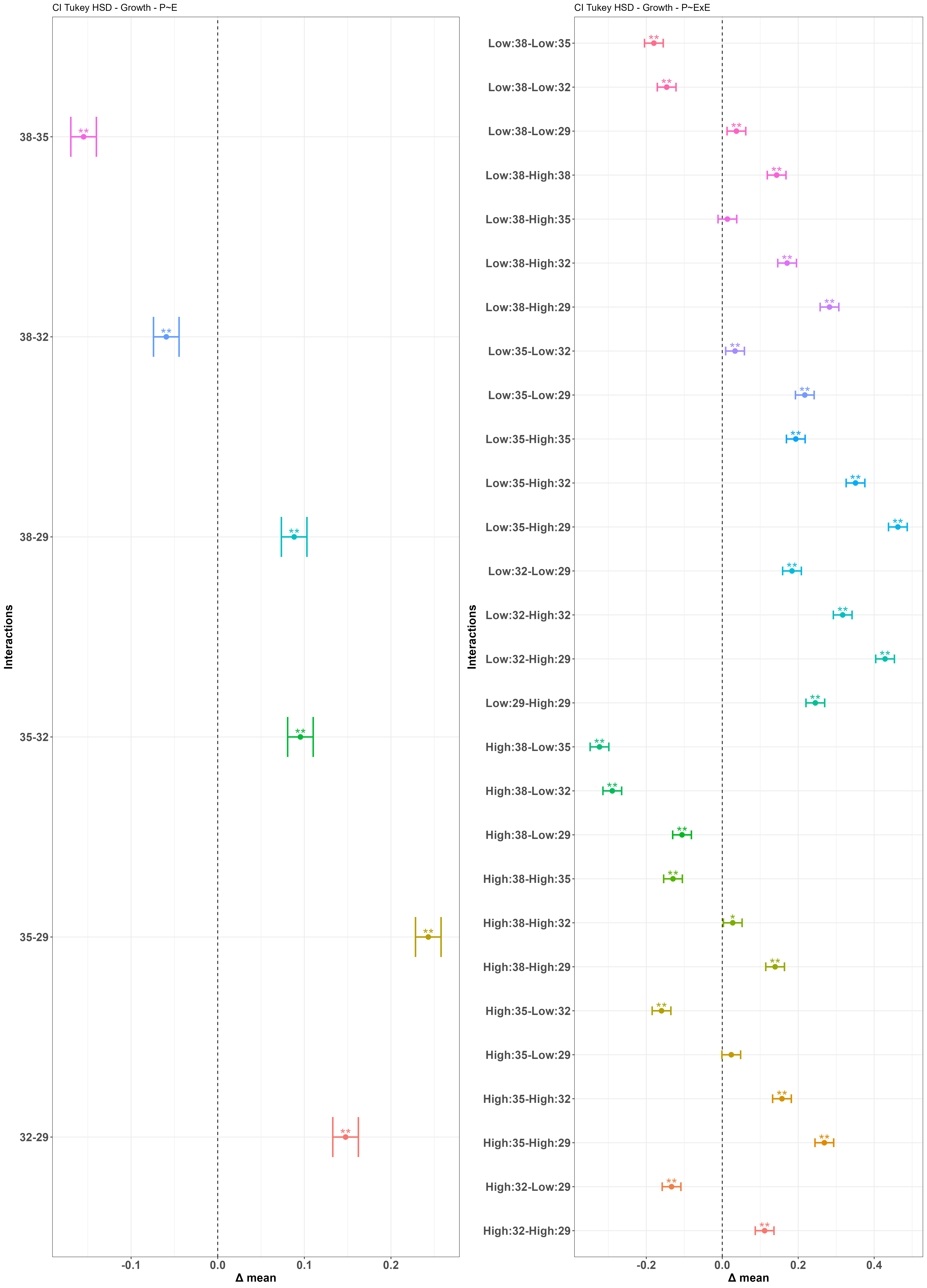

### FigS4.png

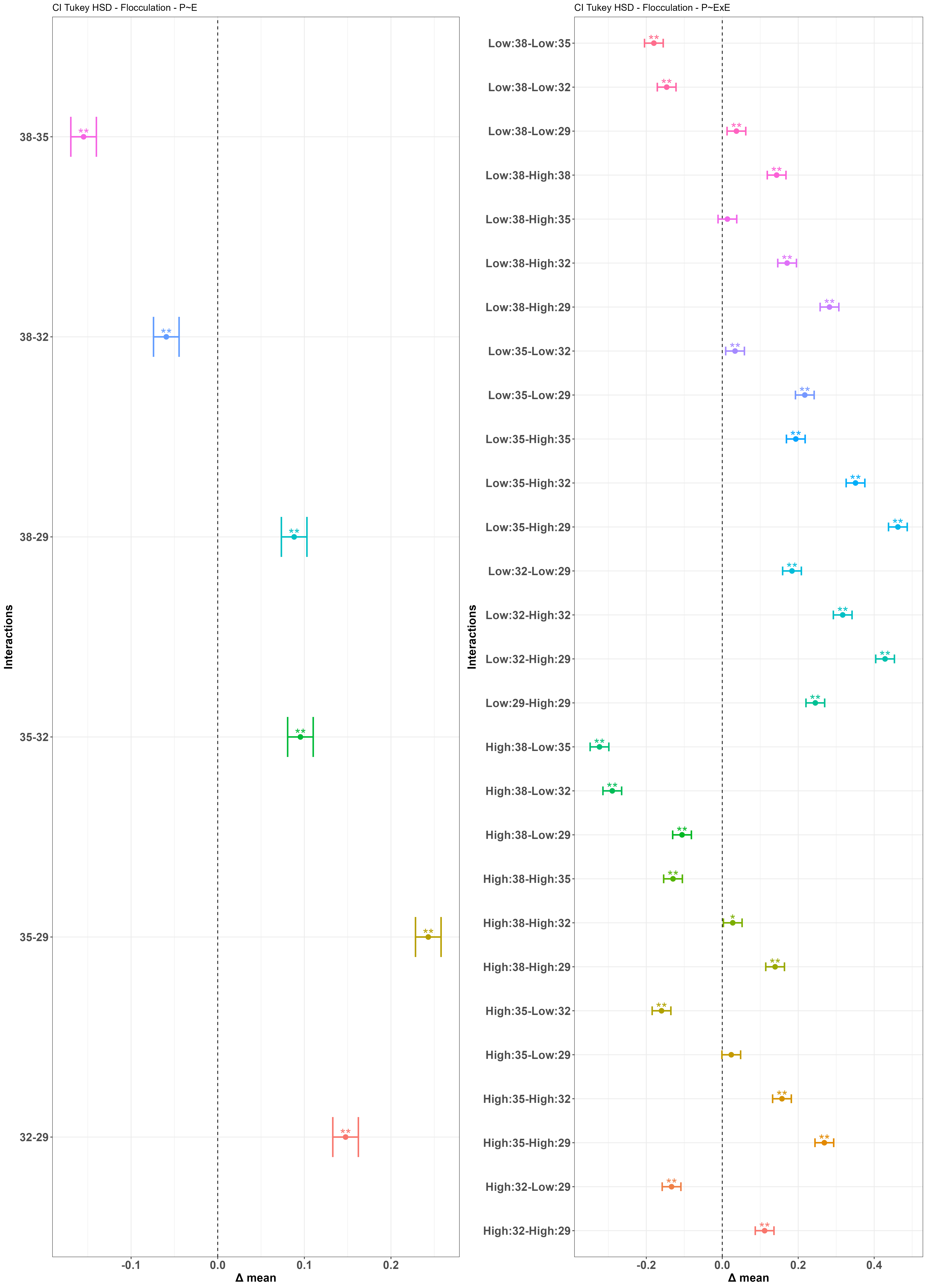

### FigS5.png

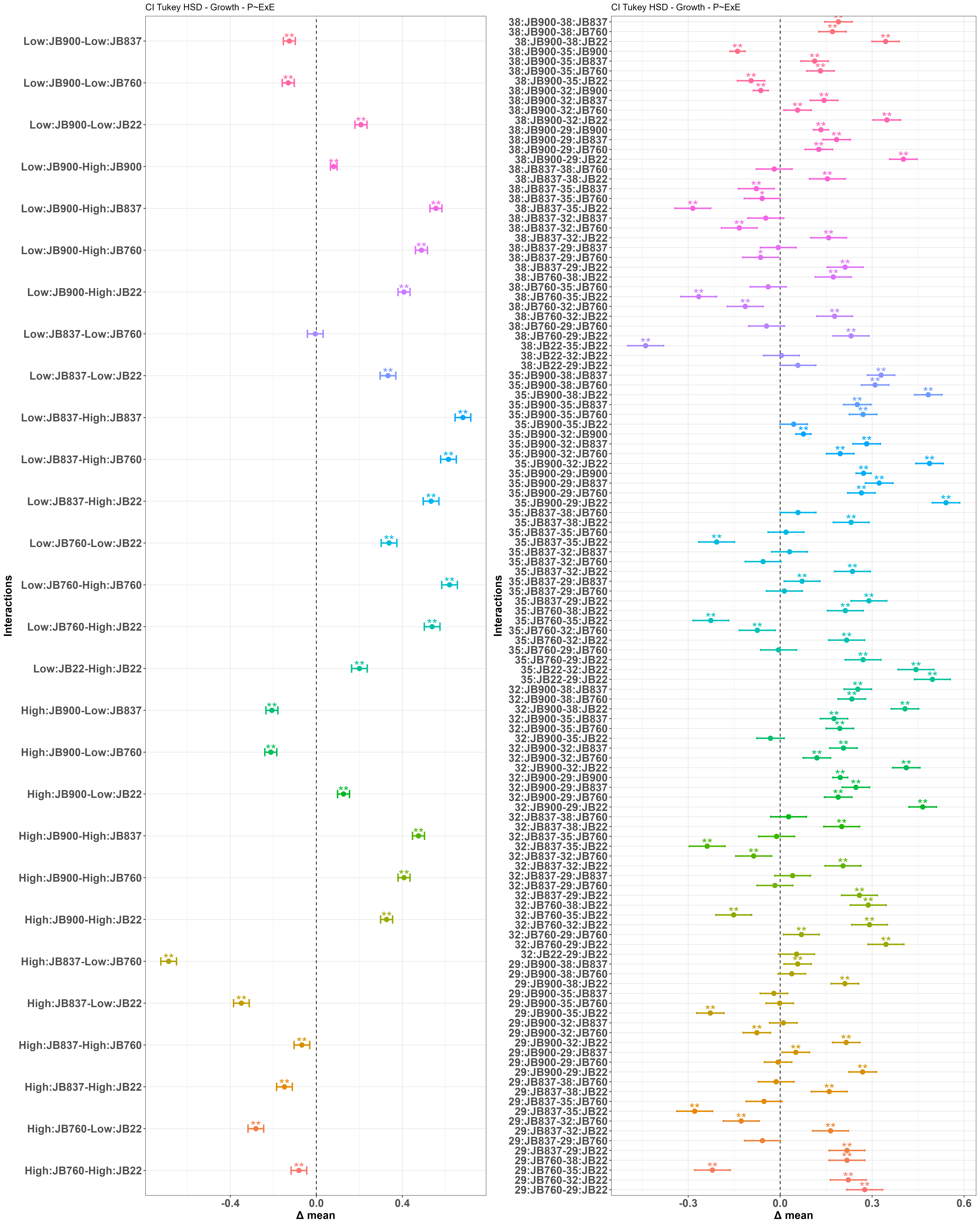

### FigS6.png

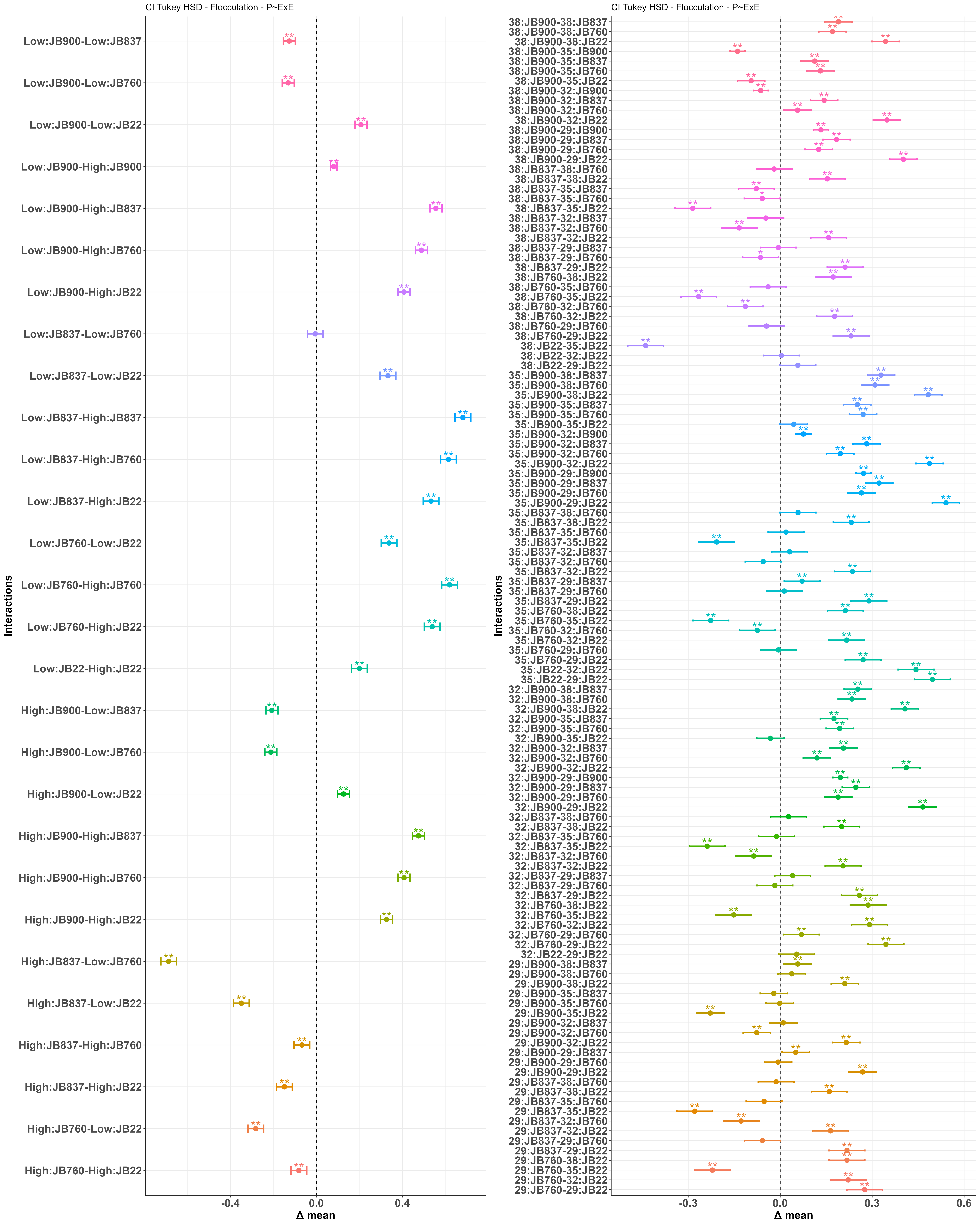
